# Pollen indicators of slash-and-burn agriculture in forest soils and peatlands: a case study of Zvenigorod biological station (Moscow region, Russia)

**DOI:** 10.1101/2025.07.14.664680

**Authors:** Ekaterina Ershova, Elena Ponomarenko, Ivan Krivokorin, Varvara Bakumenko, Nikolay Krenke, Valerii Pimenov

## Abstract

This study aimed to analyze pollen indicators of slash-and-burn agriculture in soils and aquatic deposits of the southern forest zone of Eastern Europe. Pollen spectra from buried swidden soils (500 - 1800 cal BP) were compared with coeval peat layers and modern surface soils. Swidden horizons exhibited highly distinctive pollen spectra, characterized by *Betula* dominance, low conifer pollen, and the presence of cultivation indicators (Cerealia) and post-fire succession taxa (*Chamaenerion, Pteridium, Marchantia*). The abundance of insect-pollinated *Chamaenerion angustifolium* - the most prominent indicator - suggests both its wide distribution in swidden landscapes and a possible link to ground-nesting bee activity, which may aid in identifying buried swiddens. In peat records, the onset of slash-and-burn agriculture was marked by microcharcoal, a decline in late-successional trees, a rise in pioneer taxa, and the appearance of agricultural indicators. *Betula* dominance in both archives reflects secondary stands typical of long-term swidden use. However, indicators of cultivation and succession were much less pronounced in peat. Our results show that paleosol pollen spectra provide reliable, site-specific evidence of swidden agriculture through local indicator taxa and traces of bee activity, though they tend to overrepresent local vegetation. In contrast, peat and lake records offer broader regional perspectives but detect swidden activity only indirectly. These insights provide a methodological basis for localizing ancient agricultural practices in future archeology and palaeoecology-related studies.

## Introduction

One of the most informative methods for reconstructing land use history is the palynological analysis of wetland sediments, buried soils, and occupational layers at archaeological sites. Landscape utilization has long been associated with changes in vegetation cover, including the emergence of cultivated species, the introduction of new weeds, and the expansion of ruderal flora typical of disturbed grounds. In palynology, a central approach to reconstructing land use history is the use of anthropogenic indicators—specific plant taxa in pollen spectra that reflect human activity. This method, linking particular land use types with certain pollen taxa, became widely used in Europe in the second half of the twentieth century (Iversen 1948; Behre 1981, 1986) and was later applied in many regions, including European Russia (Nosova 2009; Nosova et al. 2014; Rudenko and Novenko 2015).

Anthropogenic indicators include, first and foremost, cultivated plants, as well as taxa accidentally introduced by people from other regions. Some native species may also serve as local anthropogenic indicators if their abundance changes markedly due to human impact on vegetation.

The classical concept of pollen indicator taxa was developed primarily for interpreting pollen diagrams from lakes and peatlands—archives that accumulate airborne pollen. Certain land use types are easily recognized in such diagrams due to the presence of wind-pollinated indicator taxa. For example, pollen of *Secale* and *Fagopyrum* in wetland deposits suggests the presence of cultivated land nearby (Klerk et al. 2015; Alenius et al. 2024). Pollen of *Artemisia vulgaris*, *Chenopodium*, *Atriplex*, and *Polygonum aviculare* indicates disturbed grounds and droveways (Abraham et al. 2023). In forest zones, pollen of wild grasses, *Rumex*, *Plantago*, and other herbs are considered indicators of grazing (Alenius et al. 2020; Kamerling et al. 2021). Finally, spores of *Pteridium* and pollen of *Chamaenerion angustifolium* are associated with forest fires (Behre et al. 1986).

However, many cultivars and forage herbs are insect- or self-pollinated. Their pollen is not widely dispersed by wind and appears in aquatic sediments only if the plants grew near the water. Early cultivated plants such as *Hordeum*, *Avena*, and *Triticum* produce little airborne pollen and are poorly represented even in areas adjacent to fields (Bakels 2000). Thus, despite its power, pollen analysis of aquatic deposits often fails to address key archaeological questions, particularly those concerning specific land use types or site-level reconstructions (Hellman et al. 2009; Hjelle and Sugita 2011).

Slash-and-burn cultivation (SABC), or swidden agriculture, is based on the use of ash from in situ wood burning as fertilizer (Conklin 1957). It was practiced in Europe for millennia, until the 17th–19th centuries. According to historical sources (Petrov 1968), SABC involves forest clearing by slashing and burning trees, a short cropping phase, and a long fallow period lasting up to several decades. It was typically conducted on well-drained soils in primary coniferous or broadleaved forests; repeated cycles often occurred in secondary birch forests.

Archaeological evidence for SABC includes stored crops, the absence of tillage tools, and high settlement mobility (Lavento 2012). Although lake sediment pollen analysis lacks specific indicators of SABC, it may reflect swidden cycles through short-term antiphase fluctuations in spruce and birch pollen, along with increased charcoal concentrations (Tolonen 1978; Alenius et al. 2013; Tompson et al. 2015; Nordqvist et al. 2022). As reforestation periods shortened, birch became a constant dominant in pollen spectra (Jääts et al. 2010; Tompson et al. 2015). Pollen of cultivated plants is an unreliable indicator of early swidden agriculture, as many crops produce little (e.g., flax, rape, clover) or no pollen (e.g., root vegetables) (Erdtman 1969; Poska et al. 2018). In northern and eastern Europe, pollen of cereals appears prominently only with the spread of wind-pollinated rye in the 3th–7th centuries AD (Grikpėdis and Matuzevičiūtė 2016; Westling and Jensen 2020; Wehlin et al. 2023). Therefore, while lake pollen analysis can suggest the onset of SABC, it rarely captures individual episodes, specific crops, or the precise locations of swiddens—particularly in the areas where SABC coexisted with permanent agriculture.

Pollen analysis of soils can provide complementary information about land use, as pollen accumulation in soils at some extent differs from that in aquatic sediments (Lisitsyna et al. 2012; Geng et al. 2022). Soils lack the detailed stratification of aquatic deposits, and pollen preservation is negatively affected by good aeration and high biological activity. In addition to airborne pollen, soils receive pollen and spores from dead plant tissues, waste, dung, and manure; older pollen may also be redeposited due to erosion and lateral mass transport (Dimbleby, 1985). As a result, soil pollen spectra may differ considerably from synchronous aquatic spectra—especially in long-lived soil horizons exposed to varied anthropogenic activities and vegetation successions.

On the other hand, soils can preserve pollen and spores of local taxa that are absent from the general pollen rain and thus remain invisible in aquatic records. These include insect- and self-pollinated plants (such as many cultivars), as well as mosses and clubmosses, which produce large, heavy spores. As these are not readily wind-transported, their presence in soils may serve as a reliable site-specific indicators of past land use, including arable fields, gardens, pastures, and swiddens. Moss and clubmoss spores can accumulate in depressions through water erosion, and their concentration peaks in soil-sedimentary sequences can indicate erosional events often triggered by vegetation loss.

In recent years, we have studied soils with a documented history of agricultural use, aiming to identify indicator taxa for specific land use types in several forested and forest-steppe regions of Eastern Europe. As a result, we have described distinctive soil pollen signatures associated with dung/manure, reforested ploughlands, droveways, and waste grounds (Ershova et al. 2017; Ershova and Bakumenko 2021; Ershova et al. 2022). Our studies of forest soils in southern Estonia demonstrated that soil horizons affected by SABC in the 12th– 19th centuries exhibit diagnostic features in morphology, grain size, charcoal, phytoliths, and pollen spectra (Ponomarenko et al. 2019). Using these indicators, we identified and radiocarbon-dated more than 120 buried swidden layers across Eastern Europe (Ponomarenko et al. 2020, 2021; Krenke et al. 2022; Vyazov et al. 2019; Ponomarenko et al., in press). The pollen spectra of all these soils were highly distinctive and showed consistent features regardless of region and age.

In one of our study areas, the Zvenigorod Biological Station in the Moscow region, we are able to compare SABC-related pollen signals across multiple archives: buried swidden horizons, peat from a natural peatland, and sediments from ancient ponds. Our aim is to characterize these signals and assess their applicability for reconstruction of land use history. The new data will contribute to our knowledge of ecosystems restoration process and will be used in further archeological and palaeo-ecological studies.

## Setting

The Zvenigorod Biological Station named after C. N. Skadovsky (ZBS) is a nature preserve located in the southwestern part of the Moscow Region, approximately 40 km west of Moscow (Fig. 1). It lies within a transitional zone between the southern taiga and broad-leaved forests (Ogureeva 1999). The modern vegetation forms a mosaic of predominantly spruce, pine, and birch forests, interspersed with mixed stands of coniferous and deciduous tree species. Both boreal and nemoral elements are represented in the groundcover vegetation (Alekseev et al. 2011). Patches of floodplain meadows are preserved locally within the valleys of the Moscow River and its tributary, the Setun’ River. These meadows are currently used for hay production and, to a lesser extent, as pastures. Several large *Sphagnum* peat bogs are located on the watershed, while spring-fed fens occur on the river terraces. The soil cover consists of Podzols and Albeluvisols in automorphic settings and alluvial soils in the floodplains.

**Fig. 1.**
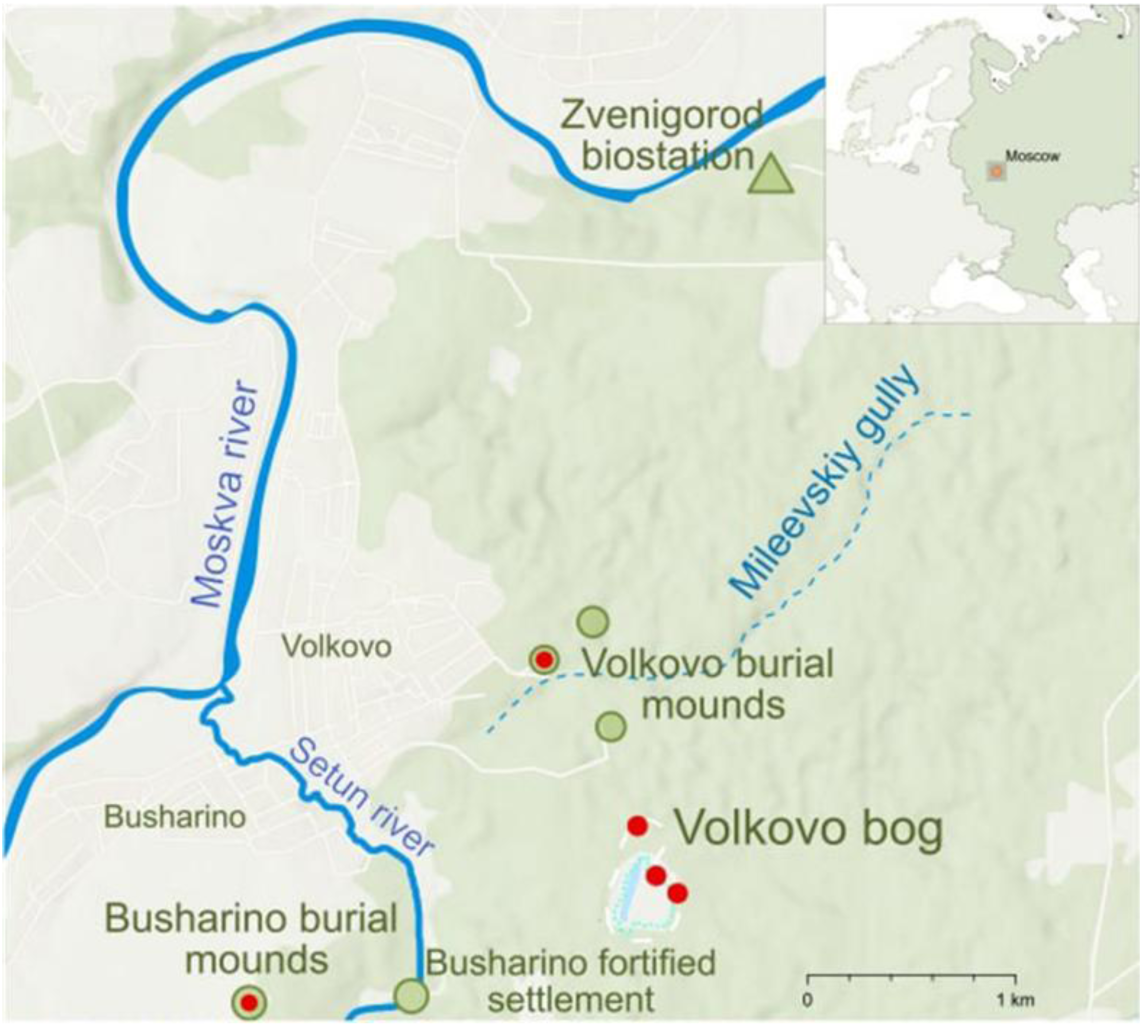
Map of the study area. Red circles indicate sampling points.

The archaeology of the area is well documented, making the ZBS an excellent model object for studying long-term land use dynamics. Archaeological studies indicate that the Moscow River region experienced multiple migration waves and several cycles of agricultural colonization from the Mesolithic to the Middle Ages (Krenke 2019). Several palynological reconstructions of land-use and vegetation dynamics in the area have been conducted using peat bogs and alluvial soils (Berezina et al. 2001; Krenke et al. 2013; Ershova et al. 2016; 2020).

Agriculture may have been practiced in the region as early as the Bronze Age (Krenke et al. 2013; Ershova et al. 2016), although this requires further investigation. Reliable evidence of agriculture first appears in deposits dating to the Early Iron Age, approximately 2800 years ago, associated with the Dyakovo culture. During the Middle Ages, agriculture became widespread due to two waves of Slavic agricultural expansion (Krenke 2019). Several necropolises with burial mounds, dated archaeologically to the 12th–13th centuries, are located within the study area (Fig. 1). According to our reconstructions and radiocarbon data, early Slavic groups inhabiting the area practiced SABC in the 10th–13th centuries (Ponomarenko et al. 2021). These same groups also constructed artificial ponds along the margins of interfluve mires for domestic purposes (Krivokorin et al. 2020).

The second wave of Slavic colonization occurred in the 14th–15th centuries, during which SABC was gradually replaced by the three-field system. This shift led to greater landscape openness. According to historical maps, by the 16th century more than 70% of the ZBS area had been cleared for settlements and farmland. Many of these ancient fields were reforested during the Time of Troubles in the 17th century (*Smuta*), while others remained in use until the late 19th century. In the 1950s, the ZBS was designated as a protected area, used exclusively for educational purposes. As a result, the current vegetation cover is largely a mosaic of former arable lands and clearcuts of various ages, now abandoned and overgrown with forest.

## Objects, methods, and chronology

### Chronology

We aimed to compare the palynological spectra of buried swidden horizons in paleosols, synchronous peat layers, Medieval pond deposits, and modern surface samples. To synchronize soil and peat layers, we radiocarbon-dated charcoal from both buried swiddens and pyrogenic peat layers. Charcoal from all pyrogenic layers was identified to the genus level, and selected fragments were submitted for AMS dating at the A.E. Lalonde Laboratory (University of Ottawa). Radiocarbon dates were calibrated using the IntCal20 calibration curve (Reimer et al., 2020). An age-depth model was constructed using the ‘rbacon’ package in the R environment (Blaauw and Christen, 2011). All radiocarbon dates used to synchronize pollen spectra, including previously published ones, are presented in Table 1.

**Table 1.**
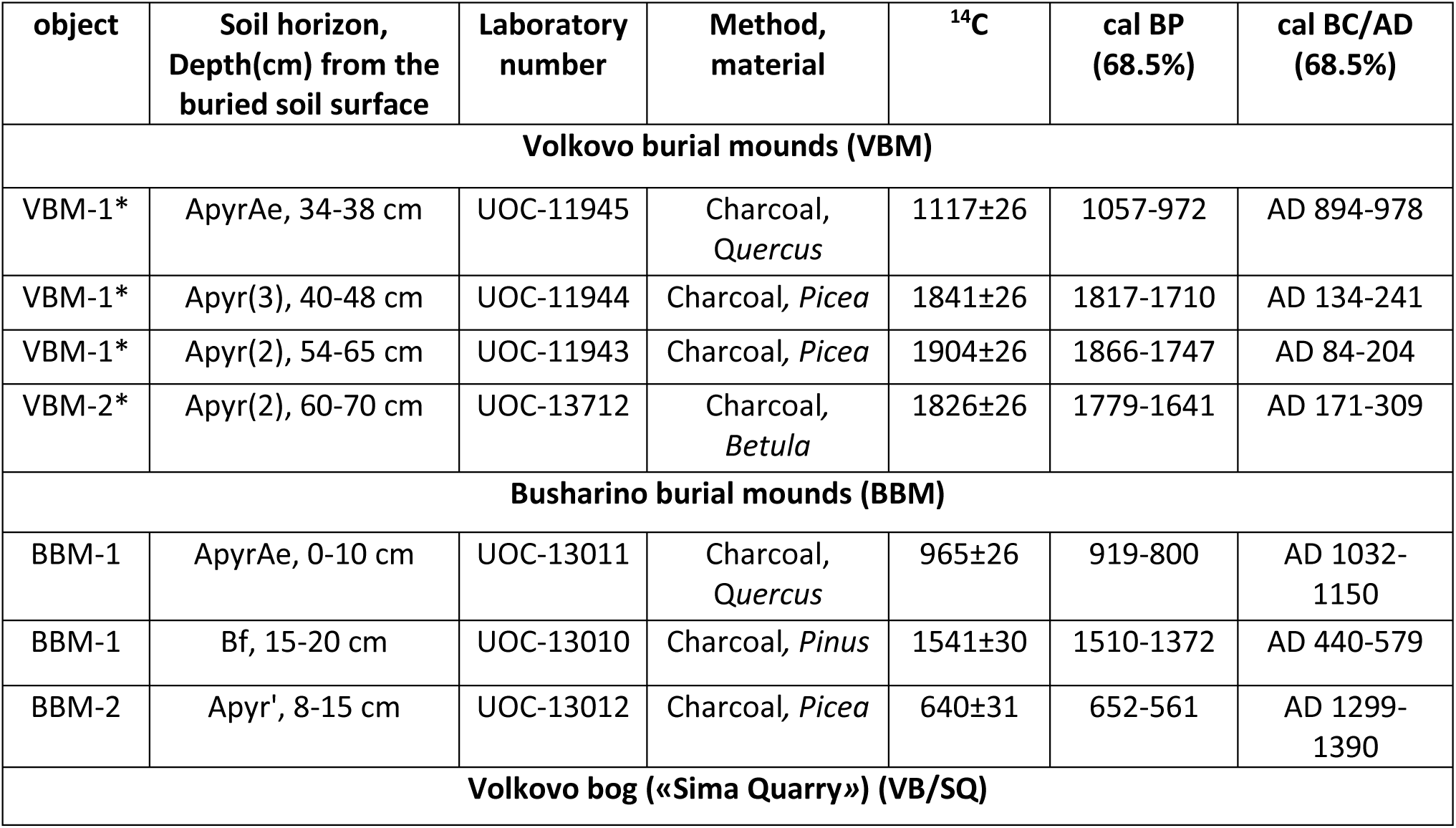
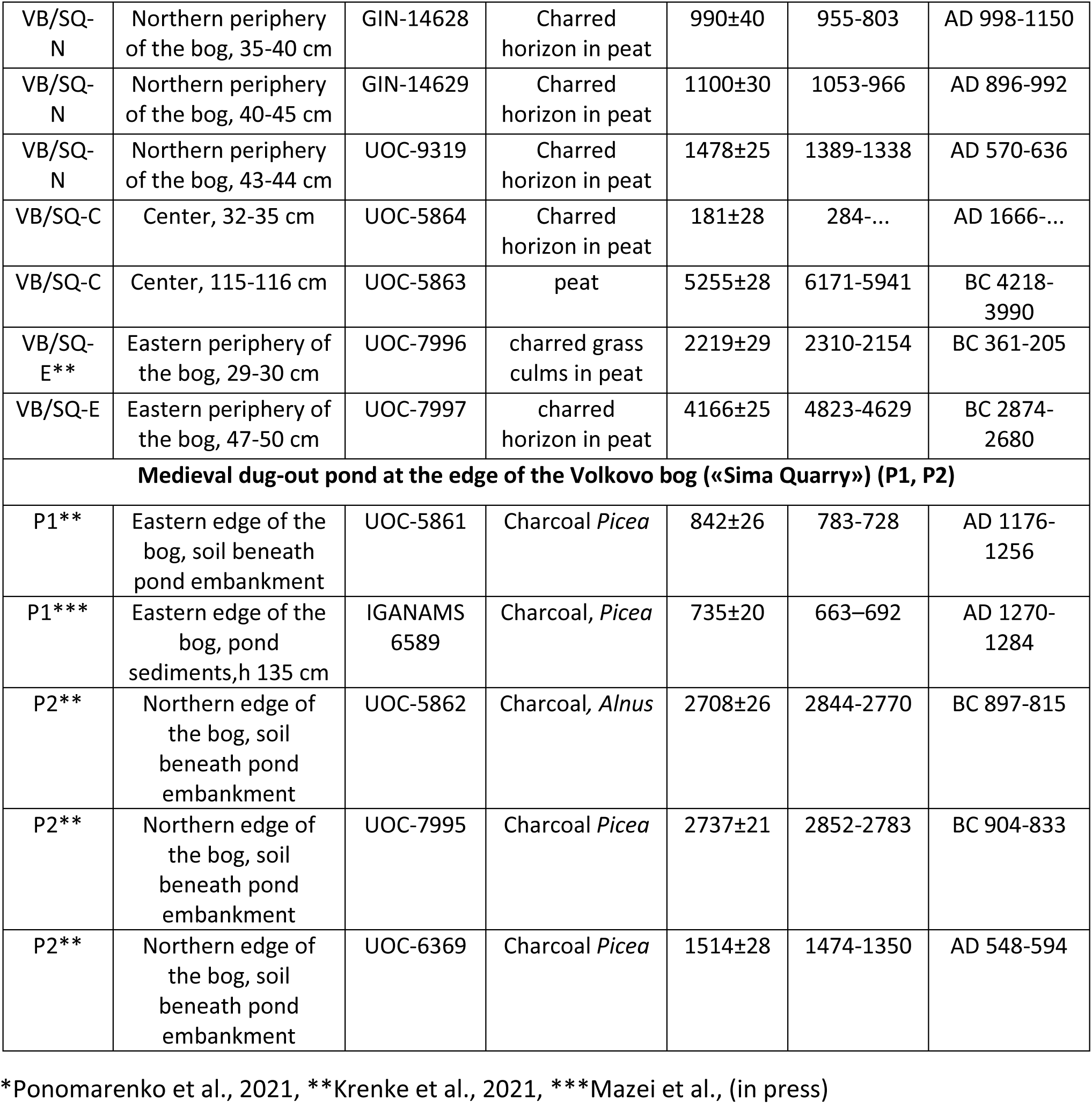
Radiocarbon dates from swidden horizons buried beneath medieval burial mounds, deposits from the Volkovo bog (“Sima Quarry”), and medieval ponds, calibrated using the IntCal20 calibration curve (Reimer et al., 2020). Asterisks indicate first published sources of the dates.

### Paleosols

To analyze traces of SABC, we sampled ancient swidden horizons buried beneath burial mounds at two medieval sites within the ZBS: the Busharino and Volkovo burial grounds, about 2.5 km apart (Fig. 1). Both sites were previously excavated and dated to the 12th–13th centuries (Krenke 2019; Krenke et al. 2020). Two burial mounds were examined at each site.

At Volkovo, three swidden layers were identified beneath one mound and two beneath the other (Fig. 2). At Busharino, one layer lay beneath a large mound, and another under slope deposits between overlapping mound toes. The buried soils at both sites showed features consistent with swidden cultivation (Ponomarenko et al. 2019): a medium gray ApyrAe horizon, 3–7 cm thick, containing infilled sweat bee burrows (∼1.5 cm in diameter) and abundant charcoal fragments (3–7 mm), with at least one >2 mm per gram of soil. Charcoal was taxonomically identified, and the most abundant taxon was selected for dating. Detailed descriptions of the Volkovo paleosols and charcoal analysis are provided in Ponomarenko et al. (2021).

**Fig. 2.**
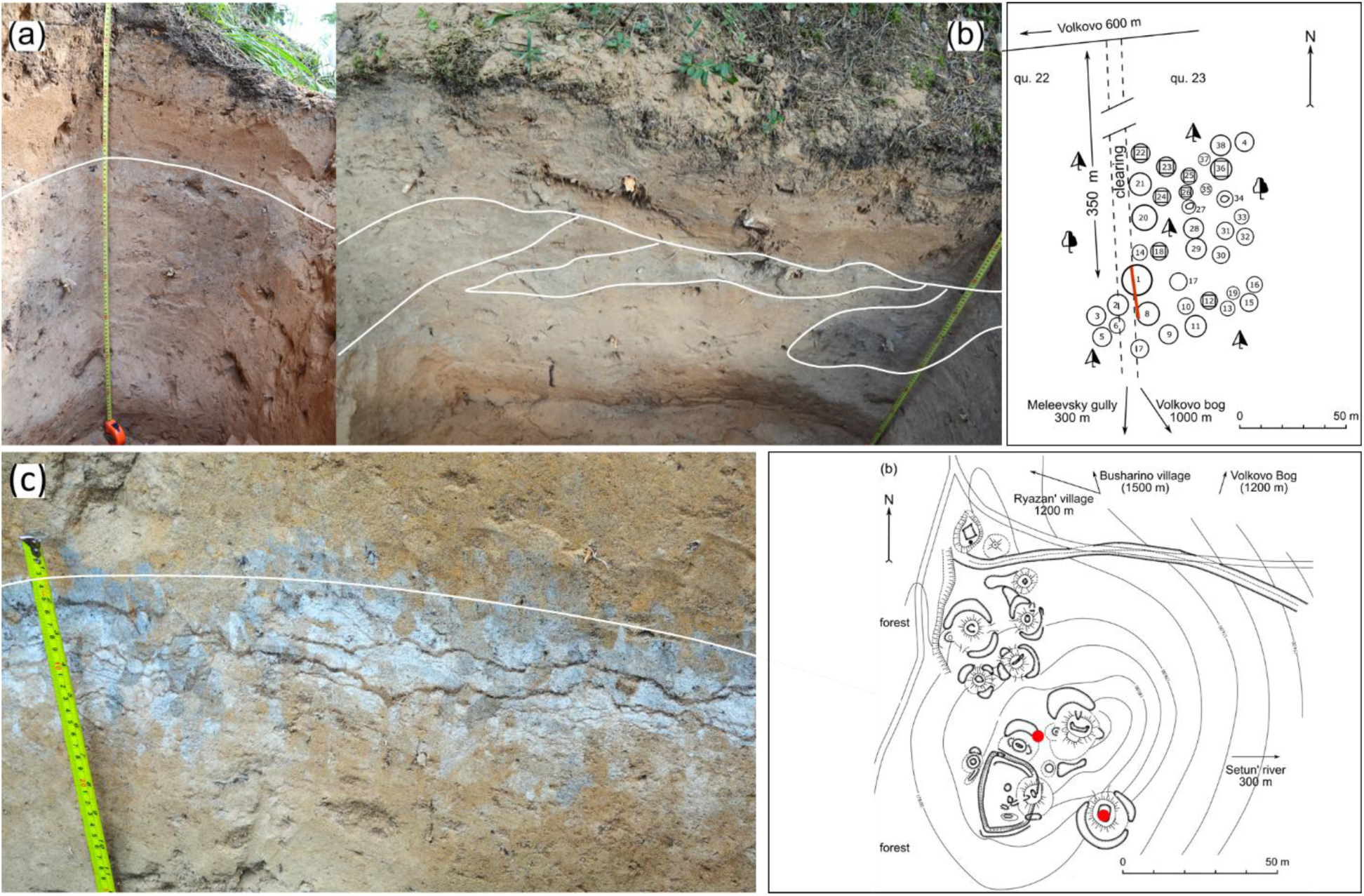
Paleosols buried beneath burial mounds (kurgans) and site plans of the Volkovo and Busharino burial grounds. Sampling points are marked with red circles. Several fragmented swidden layers, alternating with natural deposits and disturbed by uprooting, are visible beneath the Volkovo kurgans (a, b). The swidden layer beneath the Busharino kurgan lies horizontally; traces of ground-nesting bee burrows filled with charcoal-rich material are visible in the photograph (c).

**Fig. 3.**
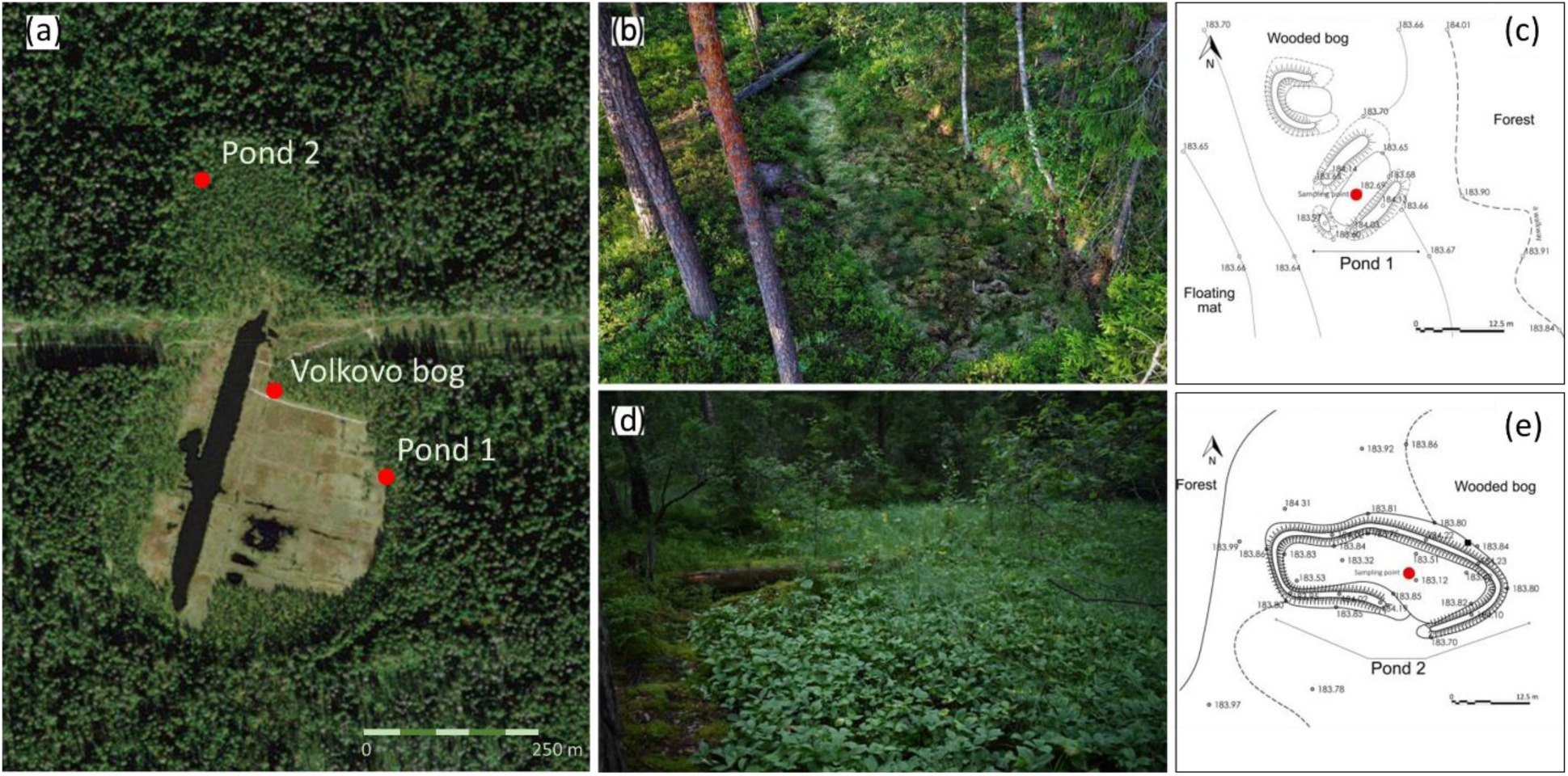
Volkovo bog (a) and medieval ponds at the edge of the bog: b, c - Pond 1, d, e - Pond 2. Sampling points are marked with red circles.

AMS dating indicates that both sites were affected by multiple SABC episodes between the 1st and 14th centuries AD (Table 1). In Volkovo, the layers date to the 1st, 2nd, and 10th centuries AD; in Busharino, to the 11th and 14th centuries AD. Thus, alongside Early Slavic swiddens that preceded mound construction, earlier swidden activity occurred during the Late Dyakovo period, and later during the second Slavic expansion.

Initial Late Dyakovo clearings targeted spruce forests, similar to kuuhta-type swiddening by Forest Finns (Holm 2007). Less than a century later, birch groves were cleared. In contrast, the Medieval population targeted oak-dominated mixed forests. After a hiatus of several centuries, swiddens of the 14th–15th centuries were again set in spruce forests. The burial mounds were constructed on former swiddens during the fallow stage—possibly years or decades after cropping ceased.

### Peatland

The Volkovo bog (“Sima Quarry”) is an ombrotrophic peatland located in a small depression on the watershed between the Moskva River and its tributary, the Setun’ River. It lies 0.5–1.7 km from the nearby Slavic burial grounds (Fig. 1) and covers ∼7 hectares. The bog’s central part was excavated during peat mining, presumably in the 18th–19th centuries, forming a water-filled depression later covered by a floating Sphagnum mat. Original peat deposits were preserved at the quarry’s margins and along the bog periphery.

Archaeological studies indicate that the peatland was part of the economic zone of nearby Dyakovo settlements between (2800–1500 cal BP) (Krenke et al. 2021), as well as of several medieval settlements during the 12th–16th centuries (Krenke, 2019).

At the peatland margins, several small ancients dug-out ponds were discovered and investigated (Krenke et al. 2021; Krivokorin et al. 2020). These features had not been previously identified at prehistoric sites. However, their location and distinct geometry suggest that peatland modification for water management was common in the past. Soils buried beneath the pond embankments contain numerous fire traces, and charcoal from these pyrogenic layers yielded radiocarbon dates corresponding to the Early Dyakovo (2840–2780 cal BP), Late Dyakovo (1474–1350 cal BP), and Early Slavic (783–728 cal BP) (Table 1).

By the 15th century, the peat bog had become part of the economic zone of a settlement located 0.5 km to the north, on the bank of the Mileyevsky Gully (Fig. 1). From the 16th to early 18th centuries, historical maps show that the bog was surrounded by fields of Volkovo village. By then, the ponds were already reintegrated into the peatland. In the second half of the 19th century, the surrounding fields were abandoned and reforested (Bakumenko and Ershova, 2021). Today, the bog and the surrounding forest form a protected area.

A 140 cm-deep peat core was extracted from the preserved part of the bog, as close to the center as possible, to capture a regional pollen signal. The basal part of the peat unit consisted of highly decomposed sedge and sedge–*Sphagnum* peat layers typical of mesotrophic conditions. The upper 40–50 cm consisted of poorly decomposed *Sphagnum* and *Eriophorum–Sphagnum* peat, characteristic of ombrotrophic bogs. Several charred layers were recorded, with the oldest dated to 4823–4629 cal BP and others spanning from 2800 to 200 cal BP (Table 1, Appendix Fig. S1).

Two ancient dug-out ponds at the eastern and northern margins of the peatland were also sampled for pollen analysis. In Pond 1, the lower 55 cm of a 150 cm core consisted of organo-mineral gyttja, with pyrogenic layers bedded at the base and at 110 cm. Radiocarbon dates—783-728 cal BP (*Picea* charcoal beneath the embankment) and 663–692 cal BP (charcoal within sediments)—place the pond in the Early Slavic colonization period, contemporaneous with nearby burial mounds. By the 15th–16th centuries, the pond was abandoned, and peat accumulation resumed; the upper 75 cm of the core consists of *Sphagnum– Eriophorum* peat.

In Pond 2, 30 cm of organo-mineral deposits were overlain by 60 cm of *Sphagnum* peat. This pond was also used by Early Slavs but was likely constructed earlier, during the Late Dyakovo period, as indicated by charcoal dated to 1474– 1350 cal BP.

### Modern surface samples

Surface samples were collected from the litter horizon in representative modern plant communities within the ZBS, including various forest types and floodplain meadows. In total, 50 samples were collected (sampling locations are detailed in Bakumenko and Ershova, 2021).

### Pollen analysis

Soil and peat samples were processed for pollen analysis using standard procedures: treatment with hot 10% KOH, separation with a heavy liquid (sodium polytungstate) for soils, followed by acetolysis (Fægri and Iversen 1989). Pollen counting was conducted under a light microscope, with 300 to 500 pollen grains counted per sample, along with additional counts of spores. Pollen diagrams were constructed using Tilia 2.0 (Grimm 1991), where stratigraphically constrained cluster analysis (CONISS) was applied on the pollen data. Community patterns were analyzed via non-metric multidimensional scaling (nMDS) in R version 4.4 (R Core Team, 2024), using the vegan package (Oksanen et al., 2016). Ordination was based on Bray-Curtis similarities of fourth-root transformed abundance data. Species significantly (p < 0.05) contributing to axis separation were visualized as bioplots using the *envfit* function.

## Results

### Surface spectra

In all modern surface spectra of ZBS, arboreal pollen (AP) contributed 85– 99% (Fig. 4). Pollen spectra of forested areas were dominated by *Pinus, Picea*, and *Betula*—the main canopy taxa. Non-arboreal pollen (NAP; 1–8%) included mainly forest herbs such as *Campanula*, *Carex (pilosa*), and Liliaceae. Anthropogenic indicators were limited to wind-pollinated *Artemisia, Chenopodium, Rumex*, and Poaceae. Polypodiaceae spores were the only type of spores represented in the samples.

**Fig. 4.**
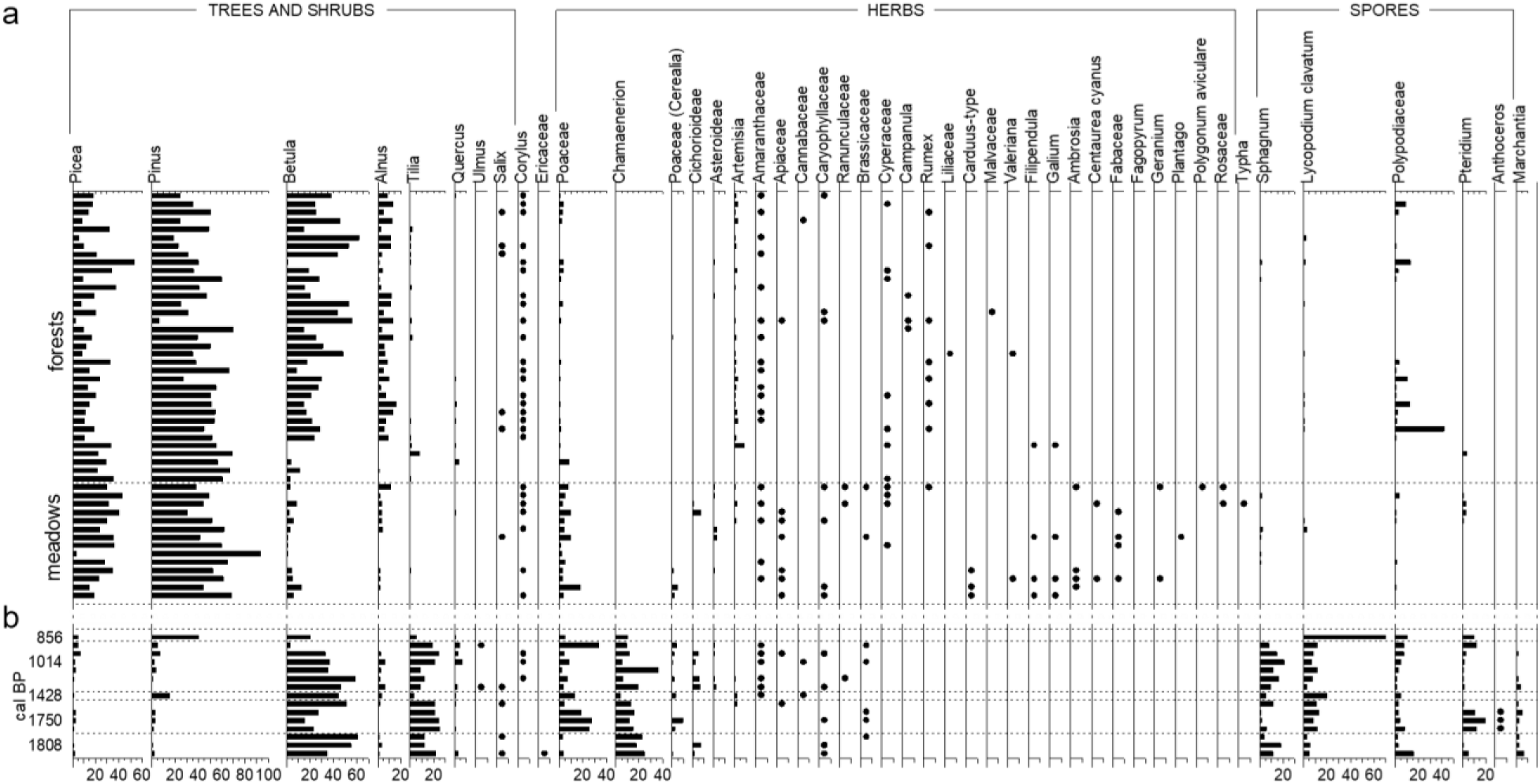
Subrecent pollen spectra from the Zvenigorod Biological Station (a) and pollen spectra from buried swidden horizons beneath burial mounds (b). Pollen taxa are shown as percentages of total pollen; spores are shown as percentages of the combined total of pollen and spores.

The meadow sites were also tree-pollen dominated, but with lower *Betula* proportions. Herbaceous pollen (up to 15%) was share higher, largely due to increased abundancy of Poaceae. The taxonomic diversity was greater due to the presence of insect-pollinated meadow taxa. Only solitary Polypodiaceae spores were found here.

### Buried swidden horizons

Pollen spectra from swidden horizons in paleosols were consistent among sites and markedly different from the modern spectra (Fig. 4). AP still dominated (60–80%) but was lower than in surface samples. *Betula* and *Tilia* prevailed, with negligible input from other trees. Herbaceous taxa were dominated by Poaceae and *Chamaenerion angustifolium*, with small amounts of Cerealia and ruderal taxa (Amaranthaceae, Cichorioideae, Caryophyllaceae, *Artemisia*). Spores were abundant—up to 70% of total sum—and included *Sphagnum*, Polypodiaceae, *Pteridium, Marchantia*, *Lycopodium clavatum*, and occasionally *Anthoceros*.

### Peatland

The pollen diagram from the Volkovo peat bog reflects vegetation changes over the past 7,000 years. According to the age–depth model, peat accumulation rates varied over time. Between 4500 and 2500 cal BP, accumulation was minimal, probably due to temporary mire desiccation. After ∼2800 cal BP, accumulation resumed but was periodically interrupted by fire events, as indicated by charred layers. Around 400–300 cal BP, the bog entered the ombrotrophic stage, and peat accumulation increased sharply. The swidden cultivation period (1800–600 cal BP), documented in soils beneath burial mounds, corresponds to a depth of 40–60 cm in the peat core (Fig. 5), where at least three charred layers are present; the basal one is dated to 2310–2154 cal BP.

**Fig. 5.**
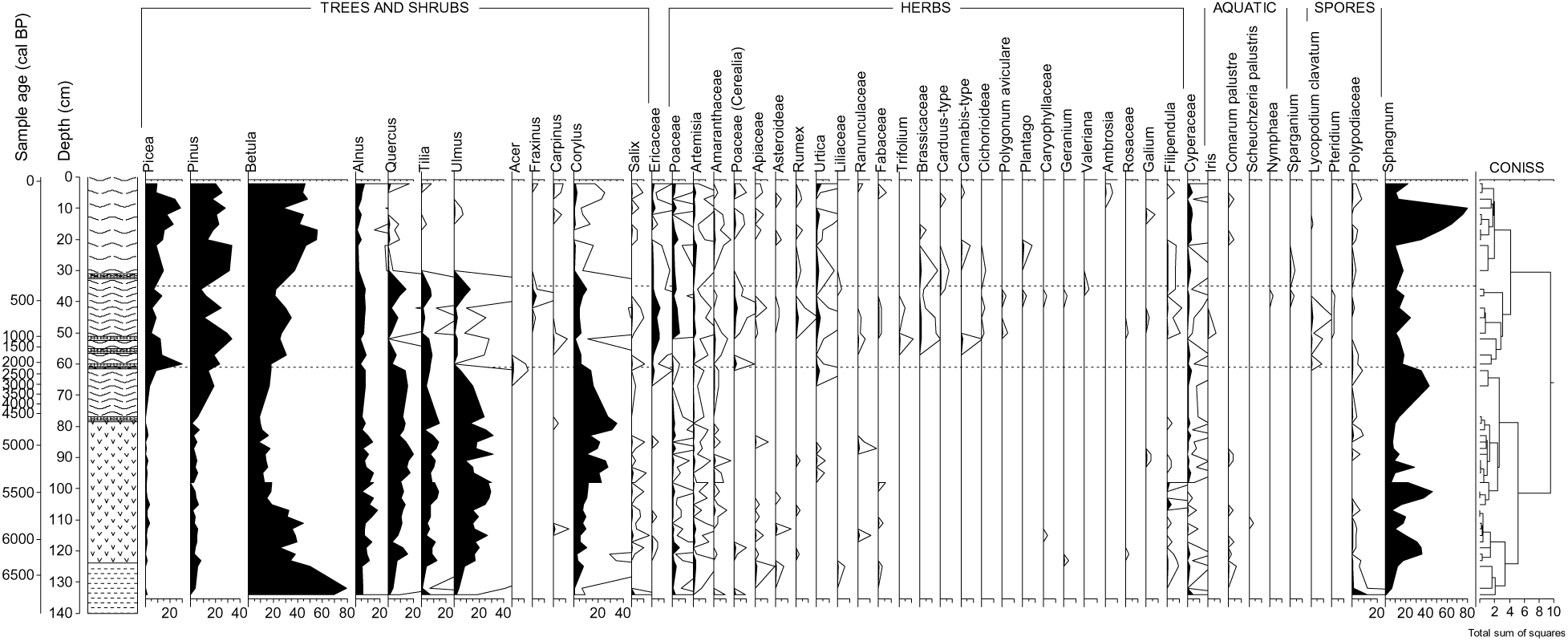
Pollen diagram of the Volkovo bog (“Sima Quarry”). Pollen taxa are expressed as percentages of total pollen; spore taxa are expressed as percentages of the combined sum of pollen and spores. The highlighted zone corresponds to the period of slash-and-burn agriculture in the ZBS area.

According to CONISS cluster analysis (Fig. 5), all peat samples corresponding to the swidden period form a distinct pollen zone. This zone begins with a sharp decline in *Picea* and broadleaved taxa, followed by an increase in *Betula*, meadow, and ruderal taxa (Poaceae, Fabaceae, Brassicaceae, Rosaceae, Apiaceae, Cichorioideae, *Urtica*, *Rumex*, *Plantago*). Cerealia-type pollen appears and peaks locally. Fluctuations are noted between primary forest taxa (*Picea, Quercus, Tilia, Ulmus*) and secondary taxa (*Betula, Alnus*). Solitary pollen grains and spores of fire and erosion indicators—*Chamaenerion angustifolium*, *Pteridium aquilinum*, *Lycopodium clavatum*—occur only in this zone. Ericaceae pollen also reaches a local maximum.

### Medieval dug-out ponds

Pollen diagrams from Ponds 1 and 2 (Fig. 6) show local vegetation dynamics over ∼800 years, offering finer resolution than the peatbog core. The lower parts (pond deposits) record land use from the 12th century until the pond’s abandonment. Pollen spectra from this period form distinct CONISS clusters (Fig. 6). *Betula* dominates, while broadleaved trees are scarce and conifers nearly absent.

**Fig. 6.**
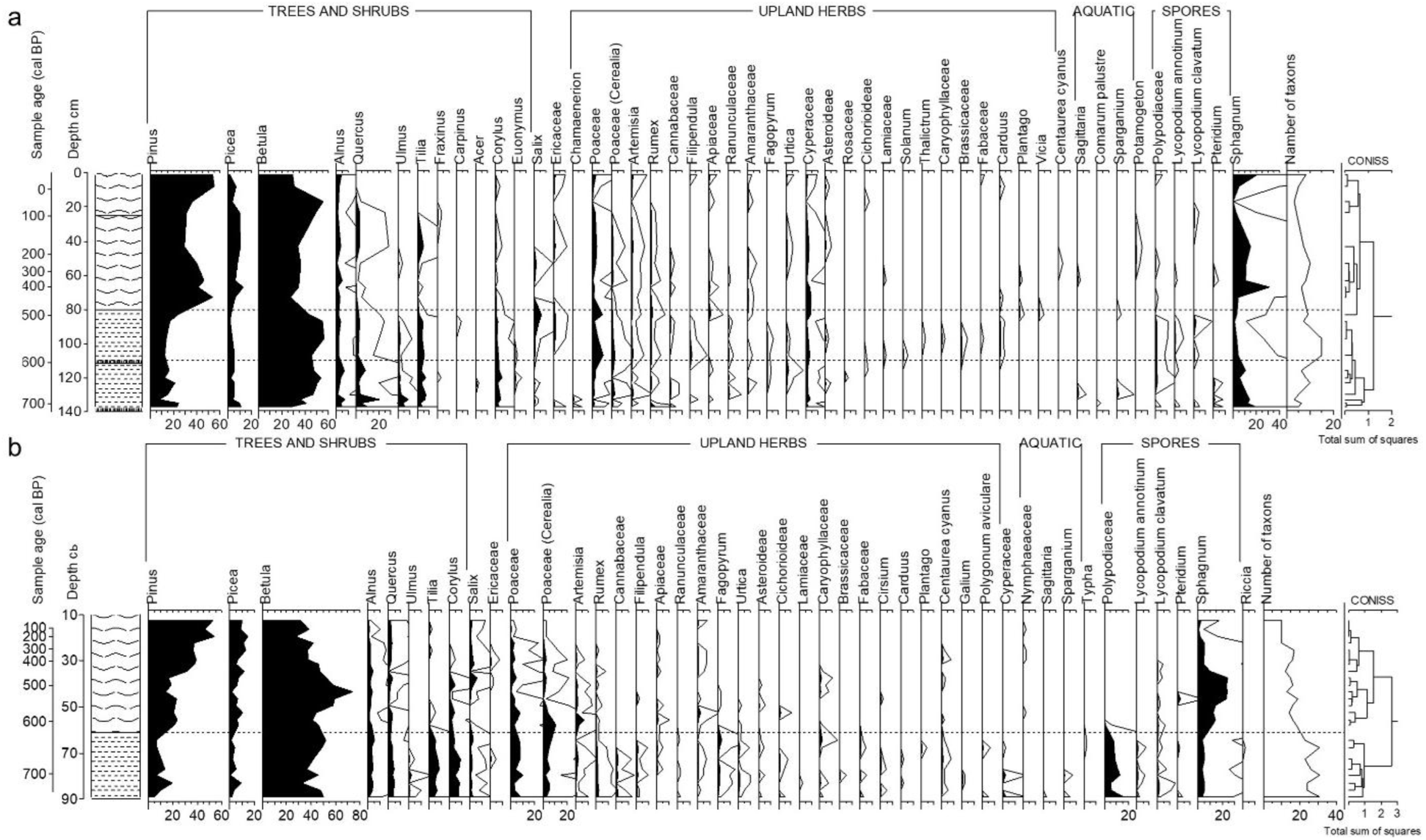
Pollen diagrams of the medieval dug-out ponds: (a) Pond 1, (b) Pond 2. Pollen taxa are expressed as percentages of total pollen; spore taxa are expressed as percentages of the combined sum of pollen and spores. The highlighted zone corresponds to the period of slash-and-burn agriculture in the ZBS area.

In Pond 1, NAP divides into two subzones. In the lower subzone, NAP is 6%, including *Secale, Fagopyrum, Artemisia, Chenopodium, Urtica*, *Rumex*, Ranunculaceae, *Filipendula*, and fire indicators (*Chamaenerion, Pteridium*). The upper subzone, above a 14th-century charcoal layer (depth: 111 cm), shows an increased NAP (17%) and a second peak in cultivated grasses. Taxonomic diversity rises, augmented by ruderal and meadow taxa (*Plantago, Carduus*, Asteraceae, Fabaceae, Caryophyllaceae, Lamiaceae, Brassicaceae, *Thalictrum*), and a peak in *Lycopodium clavatum*.

The uppermost peat layers (above 80 cm in Pond 1, 60 cm in Pond 2) differ markedly. *Pinus* pollen increases, broadleaved taxa decline, and herbaceous diversity drops. *Sphagnum* spores rise, indicating a shift to bog conditions.

## Discussion

Non-metric multidimensional scaling (nMDS) analysis (Fig. 7) illustrates the differences between the three groups of pollen spectra studied: paleosol spectra, contemporaneous peat spectra, and modern soil spectra.

**Fig. 7.**
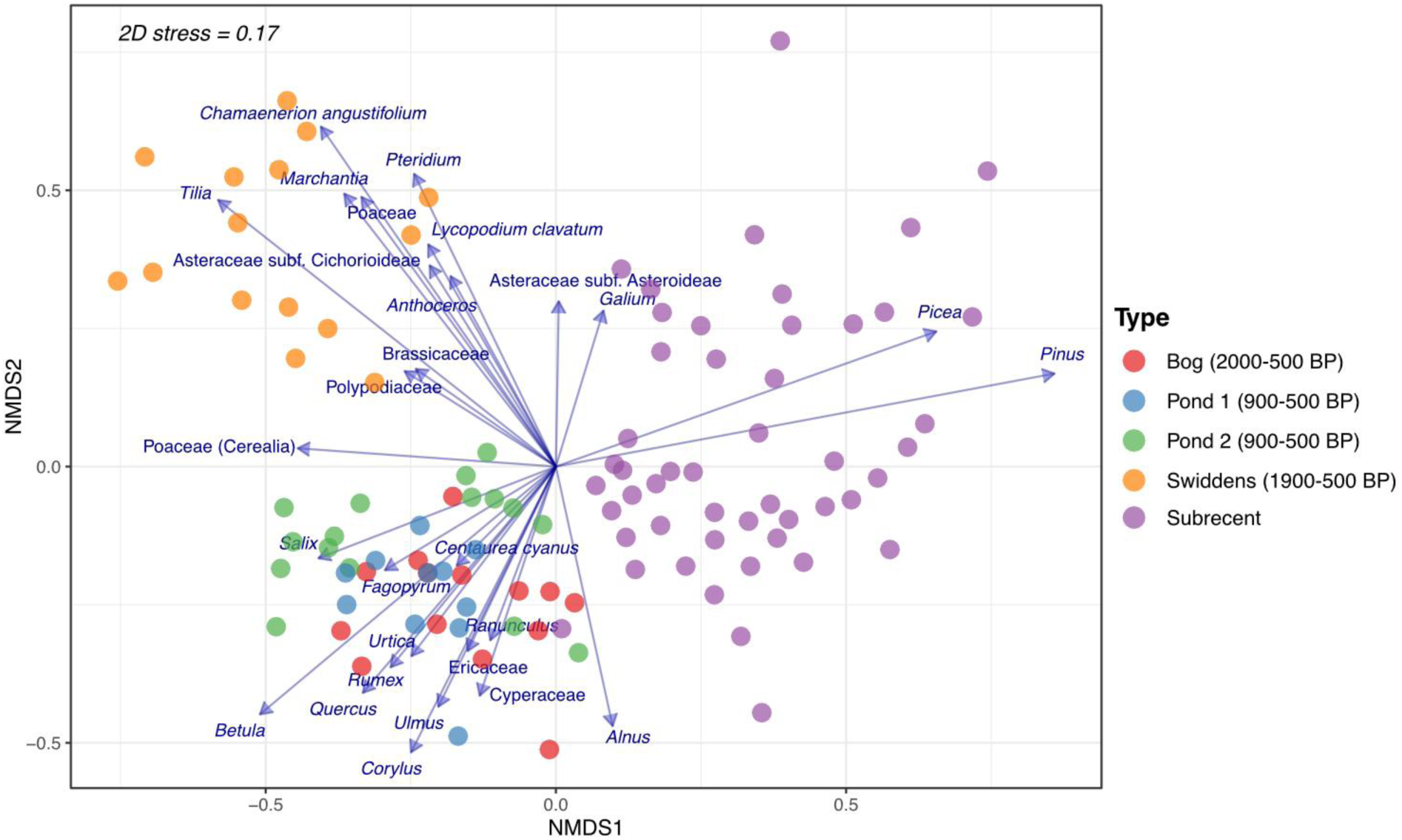
Results of non-metric multidimensional scaling (nMDS) analysis of fourth-root transformed pollen abundance data. Blue arrows indicate species significantly (p < 0.05) correlated with the ordination.

All subrecent forest pollen spectra were dominated by wind-pollinated arboreal taxa common in the modern forest cover: *Picea, Pinus*, and *Betula*, in varying proportions. This is typical for the forest zone and aligns with regional data (Nosova et al. 2020). Groundcover taxa were almost exclusively wind-pollinated herbs, comprising ≤10% of the total pollen. Meadow spectra differed slightly, showing a higher proportion of non-arboreal pollen (up to 15%) and the presence of insect-pollinated taxa. Interestingly, *Betula* pollen was underrepresented in open areas despite its local abundance and high productivity. A similar pattern was observed in the Middle Volga region (Ershova et al. 2022), suggesting that *Betula* dominance in soil spectra reliably indicates its local presence.

Pollen spectra from all buried swidden horizons (dated to 800–1800 cal BP) differed markedly. Key contrasts are highlighted by (Fig. 7): low conifer representation, dominance of *Betula* and *Tilia*, and co-occurrence of cultivation indicators (Cerealia) with post-fire succession taxa (*Chamaenerion, Pteridium, Marchantia*, *Lycopodium clavatum*). Similar spectra were observed in the Middle Volga, Upper Dnieper, and southern Estonia (Vyazov et al. 2019; Ponomarenko et al. 2020, 2019; Ershova et al. 2020).

Based on the distribution of *Betula* pollen, we suggest that its abundance reflects secondary birch groves typical of shifting cultivation landscapes. The dominance of *Betula* in swidden spectra may indicate the widespread presence of these groves. The presence of *Chamaenerion* and spores of *Lycopodium clavatum, Pteridium*, *Marchantia*, and *Anthoceros* further supports the interpretation of fire-related successional vegetation on abandoned swiddens.

In the bog and pond deposits, the swidden period is marked by charcoal layers, a decline in late-successional trees, an increase in early-successional taxa (*Betula*, *Pinus*), the appearance of cultivated and weedy taxa, and the presence of some fire indicators. In the central peat diagram, this period (2300–500 cal BP) is compressed into 22 cm (Fig. 5), but multiple clearance–regrowth episodes are still visible. The pond diagrams (Fig. 6) cover only the medieval period, capturing two waves of Slavic colonization: the earlier (10th–13th centuries), presumably associated with the Vyatichi mound-builders, and the later (14th–16th centuries), likely linked to the Muscovite population. The second wave shows a greater proportion and diversity of anthropogenic herbaceous taxa, including cultivated grasses, weeds, and pastoralism indicators.

In all diagrams—both from the central peatland and the bog-edge ponds—a distinct peak in *Lycopodium clavatum* spores and Ericaceae pollen coincides with the peaks in anthropogenic indicators. The proliferation of heather shrubs and *Lycopodium* reflects their role in successional communities on abandoned fields. Interestingly, similar co-occurrences of *Lycopodium* spores and Ericaceae pollen with anthropogenic indicators have been reported in other pollen diagrams from forest-zone bogs (Myagkaya and Ershova 2020; Novenko et al. 2016; Mazei et al. 2022).

Comparing swidden soil spectra with synchronous peat layers (Figs. 7, 8), we found common specific features:

- Both contain the same tree taxa in different proportions. In soils, *Betula* and *Tilia* dominate, while other tree species are present in minimal amounts. In peat, *Betula* is abundant but appears alongside *Pinus, Picea*, and broadleaved forest taxa.
- Non-arboreal pollen in swidden soils can reach 50%, dominated by fire indicators and ruderal taxa (*Artemisia*, Amaranthaceae, Cichorioideae, Brassicaceae, Caryophyllaceae), along with wild and cultivated grasses. In peat, NAP rarely exceeds 15% but includes a broader range: meadow taxa (Cyperaceae, *Filipendula*, *Urtica*, Lamiaceae, Fabaceae, Rosaceae) and grazing indicators (*Rumex, Polygonum aviculare*).
- Spores are more abundant and diverse in soil swiddens, while peat samples primarily contain *Sphagnum* spores.

**Fig. 8.**
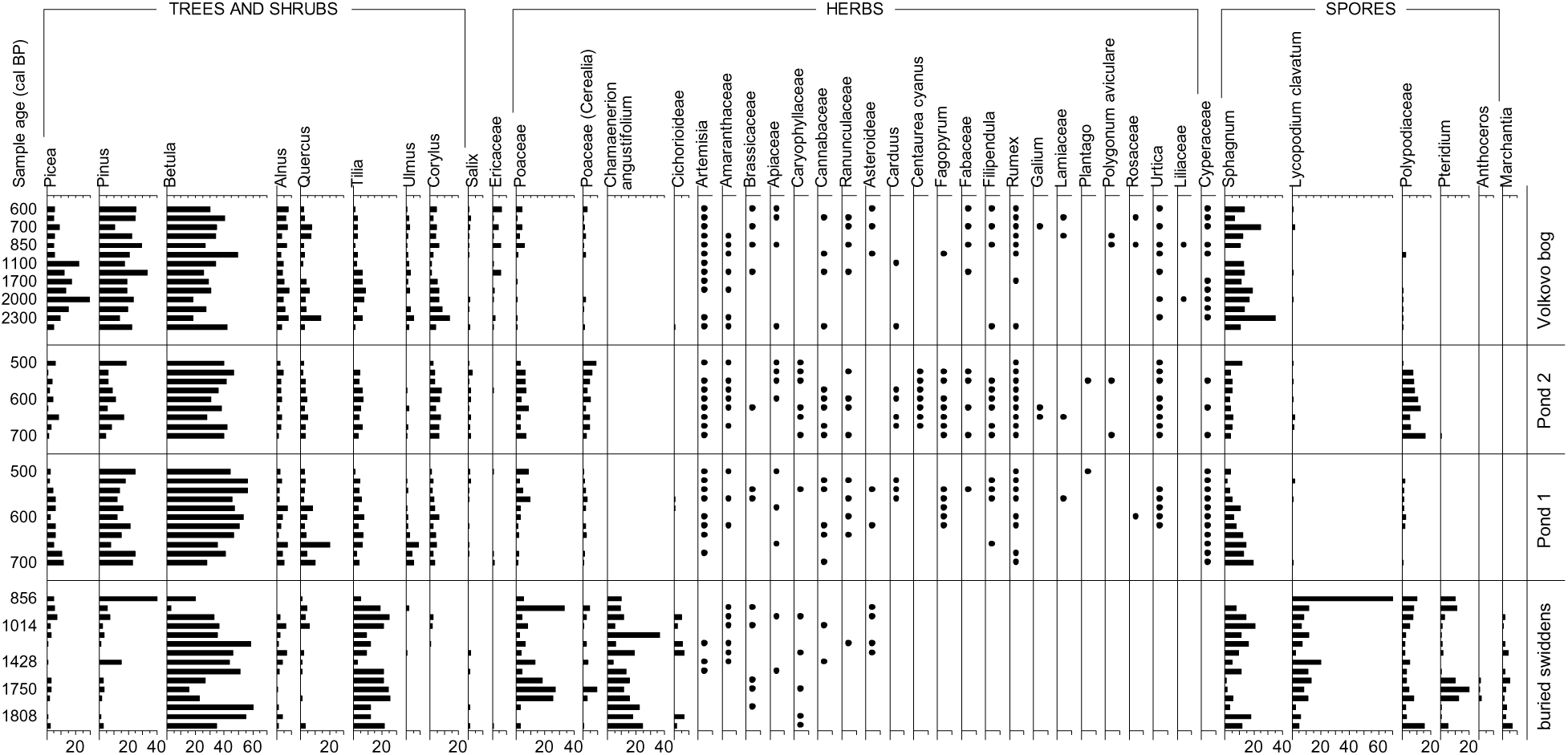
Pollen spectra of buried swidden horizons and peat tiers dating back to the period of slash-and-burn farming (2800-500 cal. BP).

We propose two main explanations of the significant differences between soil and peat swidden signatures. Firstly, soil spectra are narrowly local in both time and space, whereas peatlands larger than ∼700 m capture regional signals with finer temporal resolution. In this context, soil spectra may reflect birch groves on abandoned swiddens—hence the prominence of local taxa (*Betula, Tilia*) and fire indicators (*Chamaenerion, Pteridium, Lycopodium, Marchantia*).

Secondly, soil pollen spectra may be modified by fossorial bees, which store large amounts of pollen in their burrows, thereby distorting the pollen assemblage (Faegry, 1962; Bottema, 1975; Hunt et al 2023). All buried soils in this study showed clear evidence of bee nesting activity (Fig. 2). The exceptionally high proportion of insect-pollinated *Chamaenerion* (up to 30% of the pollen sum) may reflect this influence. Notably, bee nesting traces and high *Chamaenerion* pollen proportions are consistent features of swidden horizons across all studied sites in the southern forest zone, suggesting an important ecological role of ground-nesting bees in the early agriculture.

By the late 16th century AD, SABC had largely been replaced by permanent fields. However, prior to this, swidden and permanent agriculture may have coexisted for centuries. Our earlier research has shown that land-use transitions are clearly reflected in pollen spectra of buried soils. Regardless of regional variation, abandoned swiddens and arable fields have some distinct pollen indicators: *Chamaenerion* and *Marchantia* are diagnostic of swiddens, while *Anthoceros* and *Riccia* indicate arable fields (Bakumenko and Ershova 2021; Ponomarenko et al. 2019). These taxa are rarely found in aquatic records, as they are absent from atmospheric pollen rain.

Can SABC and arable land use be distinguished in peat records? In the central bog diagram, the two land-use types that coexisted in the Late Medieval landscape cannot be confidently separated due to low temporal resolution. However, well-stratified pond deposits (50 cm ≈ 300 years) allow clear distinction between 12th–13th-century swidden signals and 14th–15th-century tillage agriculture. The key difference lies in the abundance of herbaceous taxa associated with pastoralism— suggesting that permanent fields, established on better-drained soils, required sustained manure input. Arable farming was not feasible without sufficient livestock and pasturelands.

The massive clearance of watershed forests in the 15th–16th centuries, often referred to as the “Great Russian Clearance”, caused a major vegetation shift, recorded in all pollen diagrams from the Moscow region. This shift is marked by a sharp increase in *Betula* and *Pinus* pollen and a decline in *Picea* and broadleaved taxa to marginal levels. It is typically accompanied by increases in *Secale* and common arable weeds such as *Fagopyrum, Centaurea cyanus,* Brassicaceae, and others (Borisova 2019; Myagkaya and Ershova 2020).

## Conclusion

Our results show that identification and localization of ancient swiddens is more reliable based on the pollen spectra of paleosols than those of peatlands and waterbodies. This is due to the presence of pollen and spores from the indicator taxa that are not a part of the general “pollen rain,” but are instead deposited directly onto the soil surface by plants growing locally. Additionally, traces of ground-nesting bees—an important component of the swidden landscape— contribute to the distinctiveness of soil pollen spectra.

On the other hand, local indicator taxa cannot provide adequate information on the overall extent of swidden agriculture at the landscape or regional scale. The relative abundance of such local taxa as birch, grasses, and fireweed is greatly overestimated in soil pollen spectra compared to more regionally integrated records from peatlands and lakes.

Pollen spectra from peatlands and lakes provide a more accurate account of the proportion of agricultural land in the region and its temporal dynamics. However, they can only infer the presence of swidden agriculture indirectly, based on the presence of successional taxa, cultivated cereals, increased taxonomic richness, and elevated charcoal content. In our study, the peatland pollen diagram recorded several episodes of forest clearance followed by reforestation, with only modest increases in early-successional species following clearance events.

It appears that slash-and-burn agriculture in the ZBS area consisted of a series of local episodes, each followed by periods of depopulation and forest regrowth—at least until the onset of Slavic colonization. This allowed for the long-term coexistence of late-successional taxa (*Picea, Tilia*) and early successional taxa (*Pinus, Betula*) in the forest canopy for nearly a millennium following the introduction of forest agriculture. Complete destruction of primary forests occurred only after the 15th century, upon the large-scale clearance of watersheds for agriculture and the formation of an anthropogenic landscape.

## Acknowledgements

This work was supported by the AAUW under American Short-Term Research Publication Grants [number 018870].

## AI Usage Disclosure

Portions of this manuscript were prepared with the assistance of ChatGPT (GPT-4, OpenAI, 2024). The tool was used to improve grammar and conciseness in English-language writing. All scientific content, interpretation, and conclusions were generated and verified by the authors. No AI tools were used for data analysis, interpretation, or research design.

## Disclosure statement

The authors report there are no competing interests to declare.

## Supplemental online material

**Fig. S1.**
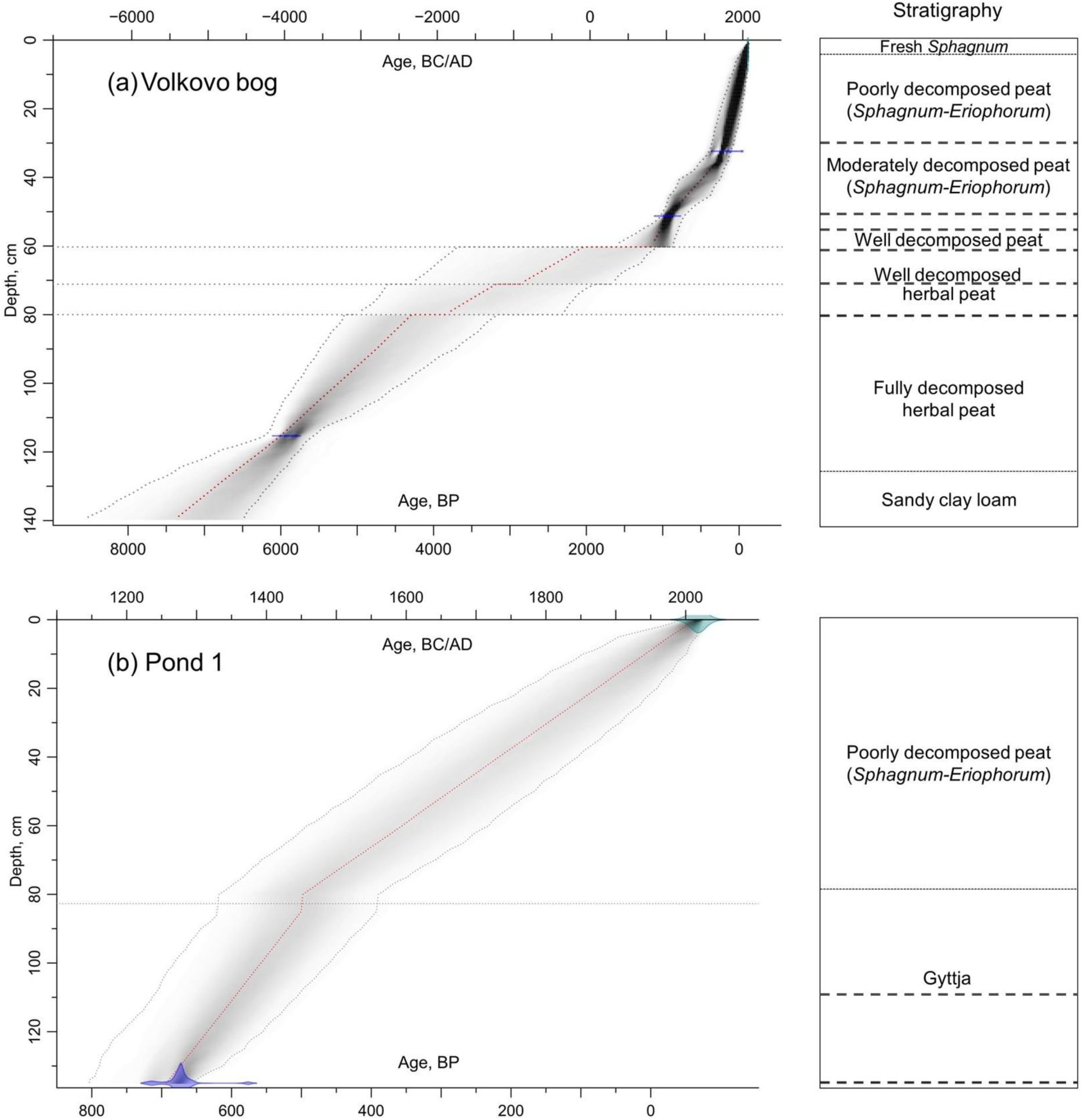
Age–depth models and core stratigraphy for Volkovo bog (a) and Pond 1 (b). Models were generated using the R package Bacon (Blaauw and Christen, 2011). The grey shaded areas represent model iterations bounded by a 95% confidence interval (dotted grey lines). In the stratigraphic diagrams, charcoal layers are marked with dashed lines.

## Notes

### Competing Interest Statement

The authors have declared no competing interest.

